# Rotavirus NSP1 contributes to intestinal viral replication, pathogenesis, and transmission

**DOI:** 10.1101/2021.06.17.448915

**Authors:** Gaopeng Hou, Qiru Zeng, Jelle Matthijnssens, Harry B. Greenberg, Siyuan Ding

## Abstract

Rotavirus (RV)-encoded non-structural protein 1 (NSP1), the product of gene segment 5, effectively antagonizes host interferon (IFN) signaling via multiple mechanisms. Recent studies with the newly established RV reverse genetics system indicate that NSP1 is not essential for the replication of simian RV SA11 strain in cell culture. However, the role of NSP1 in RV infection *in vivo* remains poorly characterized due to the limited replication of heterologous simian RVs in the suckling mouse model. Here, we used an optimized reverse genetics system and successfully recovered recombinant murine RVs with or without NSP1 expression. While the NSP1-null virus replicated comparably with the parental murine RV in IFN-deficient and IFN-competent cell lines *in vitro*, it was highly attenuated in 5-day-old wild-type suckling pups. In the absence of NSP1 expression, murine RV had significantly reduced replication in the ileum, systemic spread to mesenteric lymph nodes, fecal shedding, diarrhea occurrence, and transmission to uninoculated littermates. Of interest, the replication and pathogenesis defects of NSP1-null RV were only minimally rescued in *Stat1* knockout pups, suggesting that NSP1 facilitates RV replication in an IFN-independent manner. Our findings highlight a pivotal function of NSP1 during homologous RV infections *in vivo* and identify NSP1 as an ideal viral protein for targeted attenuation for future vaccine development.

**IMPORTANCE:** Rotavirus remains one of the most important causes of severe diarrhea and dehydration in young children worldwide. Although NSP1 is dispensable for rotavirus replication in cell culture, its exact role in virus infection *in vivo* remains unclear. In this study, we demonstrate that in the context of a fully replication-competent, pathogenic, and transmissible murine rotavirus, loss of NSP1 expression substantially attenuated virus replication in the gastrointestinal tract, diarrheal disease, and virus transmission in suckling mice. Notably, the NSP1-deficient murine rotavirus also replicated poorly in mice lacking host interferon signaling. Our data provide the first piece of evidence that NSP1 is essential for murine rotavirus replication *in vivo*, making it an attractable target for developing improved next-generation rotavirus vaccines better suited for socioeconomically disadvantaged and immunocompromised individuals.

## INTRODUCTION

Despite a dramatic reduction of rotavirus (RV) associated morbidity and mortality following the introduction of multiple safe and effective RV vaccines, group A RVs remain a major cause of life-threatening gastroenteritis among young children from 1 month to 5 years old (1, 2). RV infections still result in approximately 128,500-215,000 deaths annually worldwide (3, 4). RV vaccine option is also limited for the immunocompromised children due to the risk of persistent shedding and diarrhea (5). Thus, there remains an urgent need to develop more effective vaccines, especially for the immunosuppressed individuals.

Although RV infections in mammals occur frequently (6), RV isolates from one host species generally replicate less efficiently in heterologous species. Several RV-encoded factors, including VP3, VP4, NSP1, NSP2, and NSP3, have been implicated in contributing to this host range restriction phenotype (7). Among these viral proteins, NSP1 has been identified as an interferon (IFN) antagonist with several distinct mechanisms that enhances virus replication (8-12). NSP1 from many animal RV strains binds to and promotes the proteasomal degradation of interferon regulatory factor 3 (IRF3) (9, 13). NSP1 also recognizes and degrades IRF5, IRF7, and IRF9 (14, 15). NSP1 from several human and porcine RV strains binds to the host cullin-3 E3 ligase complex (16) and induces β-transduction repeat containing protein (β-TrCP) degradation (17). In addition, NSP1 can directly target signal transducer and activator of transcription 1 (STAT1) phosphorylation and/or translocation into the nucleus to further block the IFN amplification pathway (10, 18). Taken together, all of these studies have established NSP1 as a potent inhibitor of the host IFN responses to facilitate RV replication.

With the development of a new plasmid-based RV reverse genetics (RG) system, several groups successfully rescued recombinant simian RVs (SA11 strain), including one in which NSP1 is almost completely replaced by a NanoLuc luciferase reporter except for the first 37 amino acids at the N-terminus (19, 20). The replication of this recombinant SA11 is only modestly lower than the parental SA11 strain in MA104 cells, suggesting that NSP1 is dispensable for RV infection *in vitro*. However, since heterologous RVs replicate and spread inefficiently in mice (21), the role of NSP1 in RV infection under physiologically relevant conditions cannot be studied using this system. To overcome this hurdle, we recently constructed and recovered a fully replication-competent, infectious, and virulent recombinant murine RV using an optimized RG system (22). In this study, we take advantage of this modified RG system and further generate a new NSP1-deficient murine RV to directly address the significance and functional relevance of the NSP1 protein in intestinal replication and pathogenesis *in vivo*.

## RESULTS

### A recombinant NSP1-deficient murine RV can be successfully rescued via an optimized RG system

To determine the role of NSP1 in viral replication *in vivo*, we utilized the recombinant D6/2 murine-like RV backbone with 2 gene segments (1 and 10) derived from the simian RV SA11 strain and the other 9 gene segments (including NSP1) from the murine RV D6/2 strain (designated hereon as rD6/2-2g) (22). In order to generate an NSP1-deficient rD6/2-2g virus (rD6/2-2g-NSP1-null), we introduced two pre-mature stop codons in gene segment 5 by replacing AAG and TGC at the nucleotide positions 43 to 45, and 52 to 55 with TAG and TGA, respectively, via site-directed mutagenesis (**Fig. 1A**). With these manipulations, the protein product of gene 5 from the rD6/2-2g-NSP1-null infection is limited to the first 4 amino acids. Using the optimized RG system, we succeeded in recovering a replication-competent rD6/2-2g-NSP1-null. We next extracted the viral RNA from sucrose cushion purified RVs and performed the polyacrylamide gel electrophoresis (PAGE) to analyze the viral genomic dsRNA segments. The genomic dsRNA migration patterns were identical between rD6/2-2g and rD6/2-2g-NSP1-null viruses, with the genes 1 and 10 from SA11 and the rest 9 genes from D6/2 (**Fig. 1B**). We also validated the NSP1-null virus by a unique enzymatic digestion site (*HinfI*) introduced by the second stop codon. In comparison, cDNA amplified from gene segment 5 of rD6/2-2g was resistant to *HinfI* digestion (**Fig. 1C**). Lastly, we examined IRF3 degradation as a functional readout of murine RV NSP1 expression. To that end, we performed immunoblotting analysis to examine the cell lysates of MA104 cells infected by rD6/2-2g and rD6/2-2g-NSP1-null. With similar protein levels of RV VP6, indicating comparable replication between rD6/2-2g and rD6/2-2g-NSP1-null, the protein levels of IRF3 were undetectable in rD6/2-2g infected MA104 cells whereas IRF3 was not degraded by rD6/2-2g-NSP1-null infection (**Fig. 1D**). Based on these results, we demonstrate that the rD6/2-2g-NSP1-null virus was successfully rescued and did not express the NSP1 protein.

**Figure 1.**
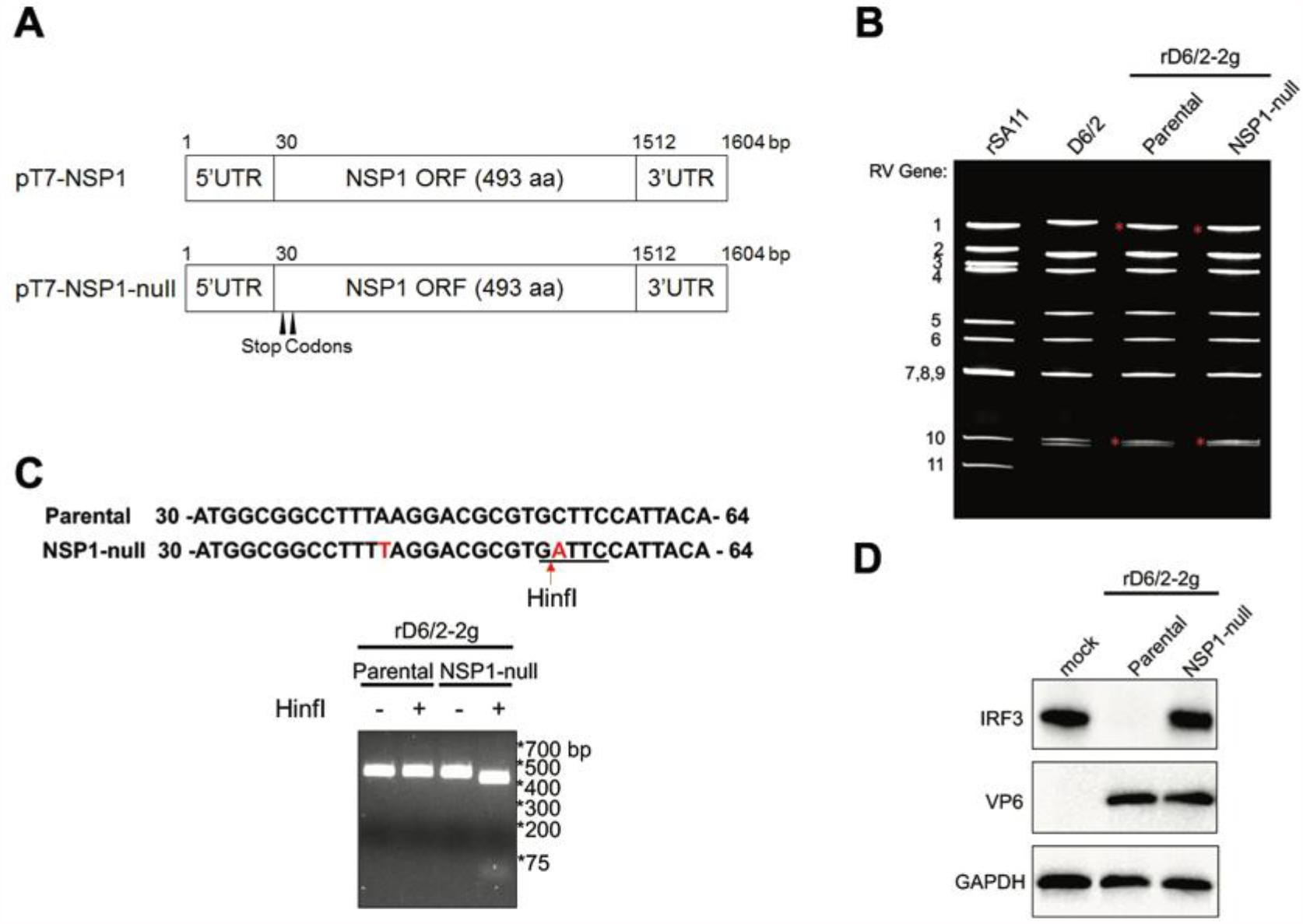
Generation of a recombinant NSP1-deficient murine RV using the optimized reverse genetics system. (**A**) Schematics of the plasmids used to rescue rD6/2-2g (pT7-NSP1) and NSP1-deficient rD6/2-2g-NSP1-null (pT7-NSP1-null) viruses. To generate the rD6/2-2g-NSP1-null, AAG and TGC at the nucleotide positions 43 to 45, and 52 to 55 were replaced with two stop codons TAG and TGA, which are indicated by the black arrowheads. UTR, untranslated region; ORF, open reading frame; bp, base pairs; aa, amino acids. (**B**) RNA was extracted from sucrose gradient concentrated indicated RV strains, separated on a 4-15% polyacrylamide gel, and stained by ethidium bromide. Genes 1 and 10 from SA11 strain were marked by red asterisks. (**C**) The stop codon introduced at nucleotides position 52 to 55 creates a unique *HinfI* digestion site. The NSP1 5’ end PCR products were digested by *HinfI* at 37°C for 1 h and separated by a 2% agarose gel. (**D**) MA104 cells were infected by parental rD6/2-2g or rD6/2-2g-NSP1-null at an MOI of 3 for 6 h. The infected cells were lysed by RIPA buffer and the protein levels of IRF3, VP6, and GAPDH in the cell lysates were analyzed by immunoblotting using indicated antibodies.

### The replication of a recombinant NSP1-deficient murine RV is comparable to the parental murine RV in multiple cell lines

To determine whether the loss of NSP1 protein negatively impacts virus replication in cell culture, we performed a multi-step growth curve for rD6/2-2g and rD6/2-2g-NSP1-null in MA104 cells at a multiplicity of infection (MOI) of 0.01. Both focus-forming unit (FFU) and quantitative reverse transcription-polymerase chain reaction (RT-qPCR) assays did not reveal significant differences between rD6/2-2g and rD6/2-2g-NSP1-null over the time course (**Fig. 2A and B**). The plaque sizes of rD6/2-2g-NSP1-null (diameter, 2.07 ± 0.53 mm) were significantly smaller than those of rD6/2-2g (diameter, 4.53 ± 0.99 mm) in MA104 cells (**Fig. 2C**). We also validated the genetic stability of rD6/2-2g-NSP1-null by serially passaging the virus for 5 times in MA104 cells and confirming the presence of two stop codons in gene segment 5 (**Fig. 2D**). These results are consistent with the previous report that an intact NSP1 is not required for simian RV SA11 strain infection in MA104 cells (19, 20).

**Figure 2.**
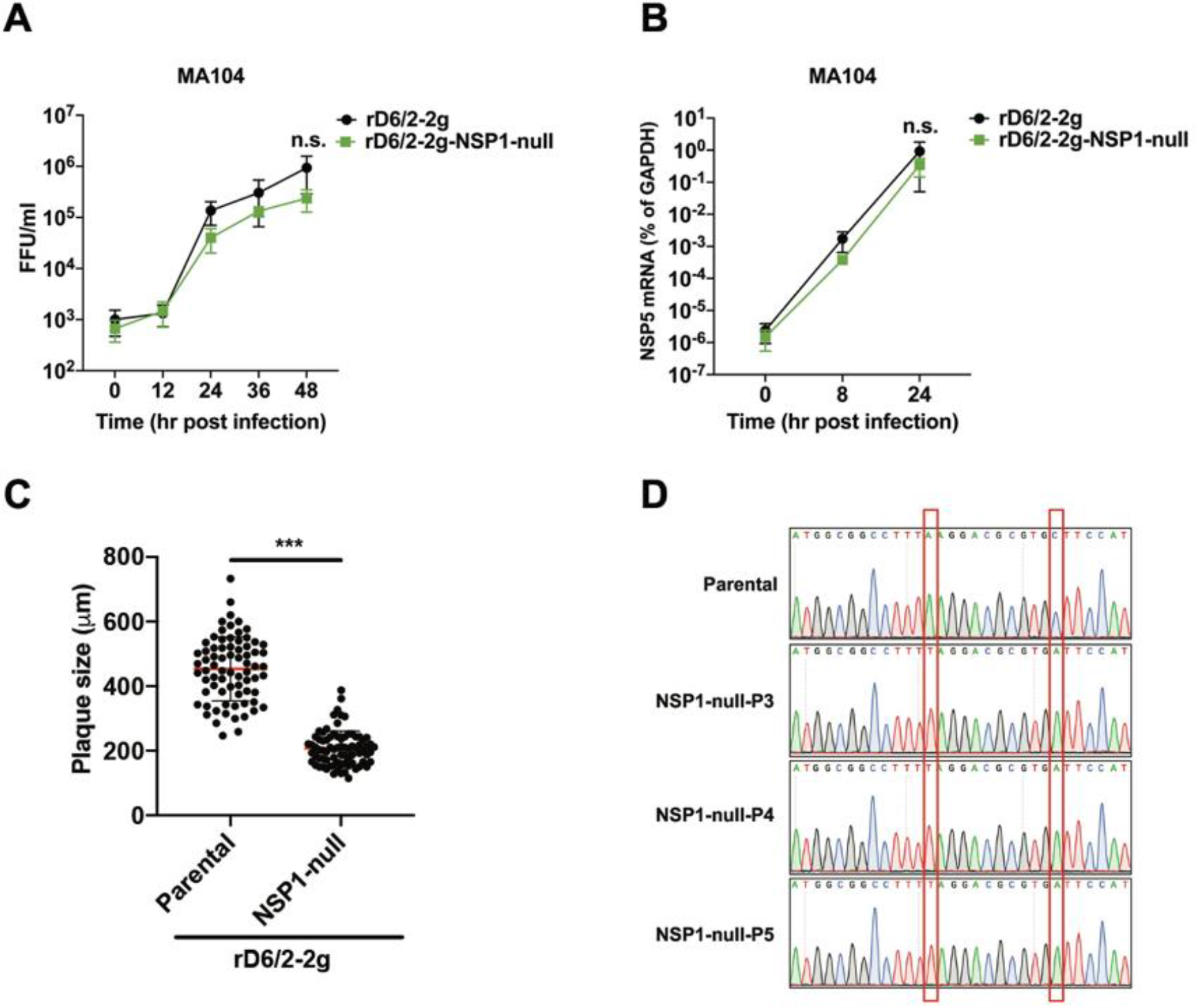
Replication kinetics of rD6/2-2g and rD6/2-2g-NSP1-null in MA104 cells. (**A**) MA104 cells were infected with rD6/2-2g or rD6/2-2g-NSP1-null at an MOI of 0.01 (FFU/cell) for 0, 12, 24, 36, and 48 h. Infected cells were harvested by freezing and thawing for 3 times. The titer of the propagated viruses at different time points were determined by an FFU assay. The data displayed were the mean ± SD for three different assays. (**B**) MA104 cells were infected as described above for 0, 8, and 24 h. RNA was extracted from infected cells and RT-qPCR was used to measure RV NSP5 transcript levels in infected cells. The data shown were the mean ± SD for three different assays. (**C**) Plaque assays were performed for rD6/2-2g or rD6/2-2g-NSP1-null in MA104 cells at an MOI of 0.01 and individual plaque formation was recorded and measured at 5 dpi by a bright-field microscope. The data shown are the mean ± SD for two different assays. (**D**) Indicated recombinant murine RVs were serially passaged 5 times in MA104 cells. The NSP1 fragments were amplified from the viruses and analyzed by Sanger sequencing. *** *P*<0.001; n.s., not significant (unpaired student’s *t* test).

Since NSP1 dampens the host IFN responses, which are defective in MA104 cells, we next tested whether the replication of rD6/2-2g-NSP1-null is restricted in IFN-competent cell lines. We examined the growth kinetics of rD6/2-2g and rD6/2-2g-NSP1-null in two different cell types: HEK293 and HAP1 cells, which are human embryonic fibroblastic cell line and human myeloid leukemia cell line, respectively. Although these cells mount robust type I and III IFN responses to RV infection (23), rD6/2-2g-NSP1-null still replicated comparably to rD6/2-2g in these IFN-competent cell lines (**Fig. 3A and B**), suggesting that NSP1 is dispensable for murine RV replication in a cell-type independent manner.

**Figure 3.**
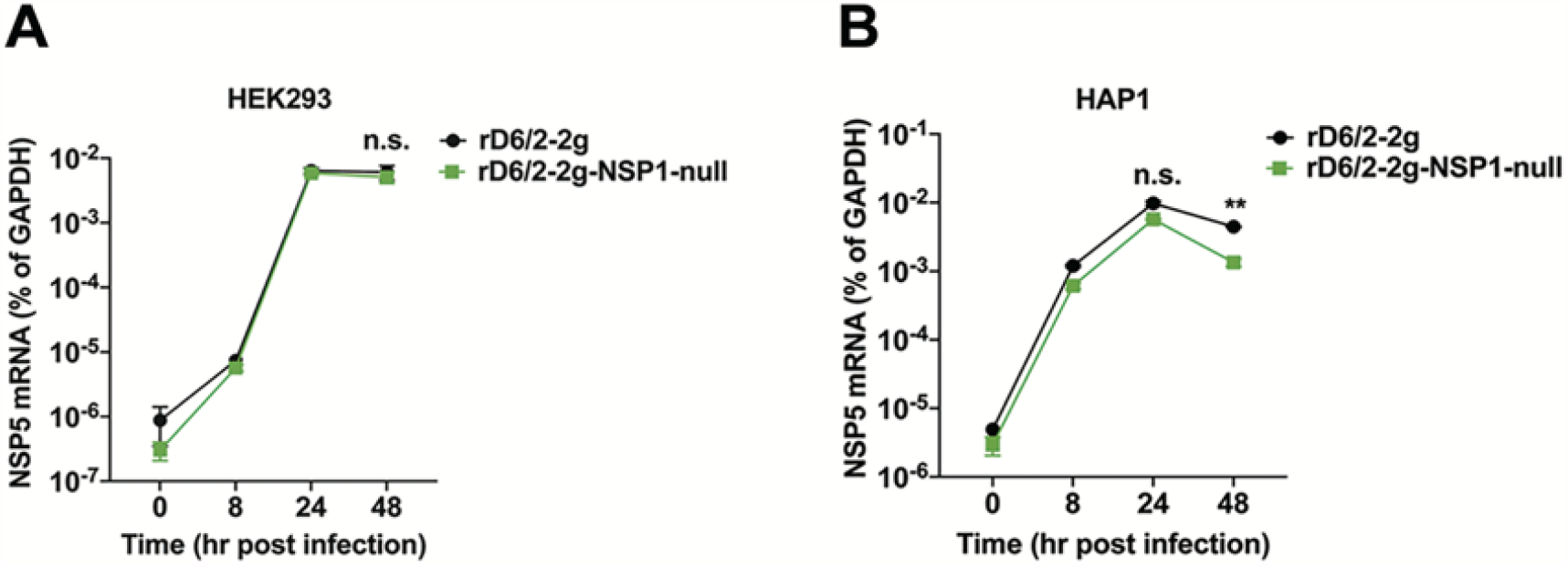
Growth curves of rD6/2-2g and rD6/2-2g-NSP1-null in IFN-competent cells. HEK293 (**A**) and HAP1 (**B**) cells were infected by rD6/2-2g and rD6/2-2g-NSP1-null at an MOI of 0.01 for 0, 8, 24, and 48 h. The expression level of NSP5 was quantified by RT-qPCR. The data shown are the mean ± SD for three individual assays. ** *P*<0.01; n.s., not significant (unpaired student’s *t* test).

### Loss of NSP1 severely attenuates murine RV replication in 129sv suckling mice

To compare the replication, pathogenesis, and spread of rD6/2-2g and rD6/2-2g-NSP1-null in an *in vivo* environment, we orally inoculated 5-day-old wild-type 129sv suckling mice with 1.5×10^3^ FFUs of rD6/2-2g or rD6/2-2g-NSP1-null and monitored the diarrheal development from day 1 to 12 post infection. While the diarrhea occurrence from the rD6/2-2g infected group was consistently higher than 70% for the first 10 days, rD6/2-2g-NSP1-null caused minimal to no diarrhea in infected animals for the first 2 days (**Fig. 4A**). Starting from 3 days post infection (dpi), rD6/2-2g-NSP1-null started to approximate the parental virus, with their curves eventually trending in a similar fashion (**Fig. 4A**). We also quantified the shedding of infectious RVs in the feces of mouse pups by an FFU assay. Consistent with the defects in diarrhea, fecal shedding of rD6/2-2g-NSP1-null in infected mice could not be detected at 1 and 2 dpi, whereas we observed high levels of RV titers from the rD6/2-2g infection (**Fig. 4B**). The two virus shedding curves looked similar from 3 to 5 dpi, before the RV shedding in rD6/2-2g-NSP1-null infected pups waned again as compared to that of rD6/2-2g group (**Fig. 4B**). Furthermore, we also evaluated the ability of these two viruses to transmit to uninfected littermates, an important trait for viruses that spread fecal-orally. We found that virtually all the non-inoculated littermates of the rD6/2-2g infected pups developed diarrhea at 6 dpi (**Fig. 4C**). In comparison, the maximal percentage of diarrhea among mock infected pups in the rD6/2-2g-NSP1-null cage reached 40% and lasted only 1 day (**Fig. 4C**). Collectively, these results suggest that NSP1 is necessary for optimal RV infection, disease, and spread *in vivo*.

**Figure 4.**
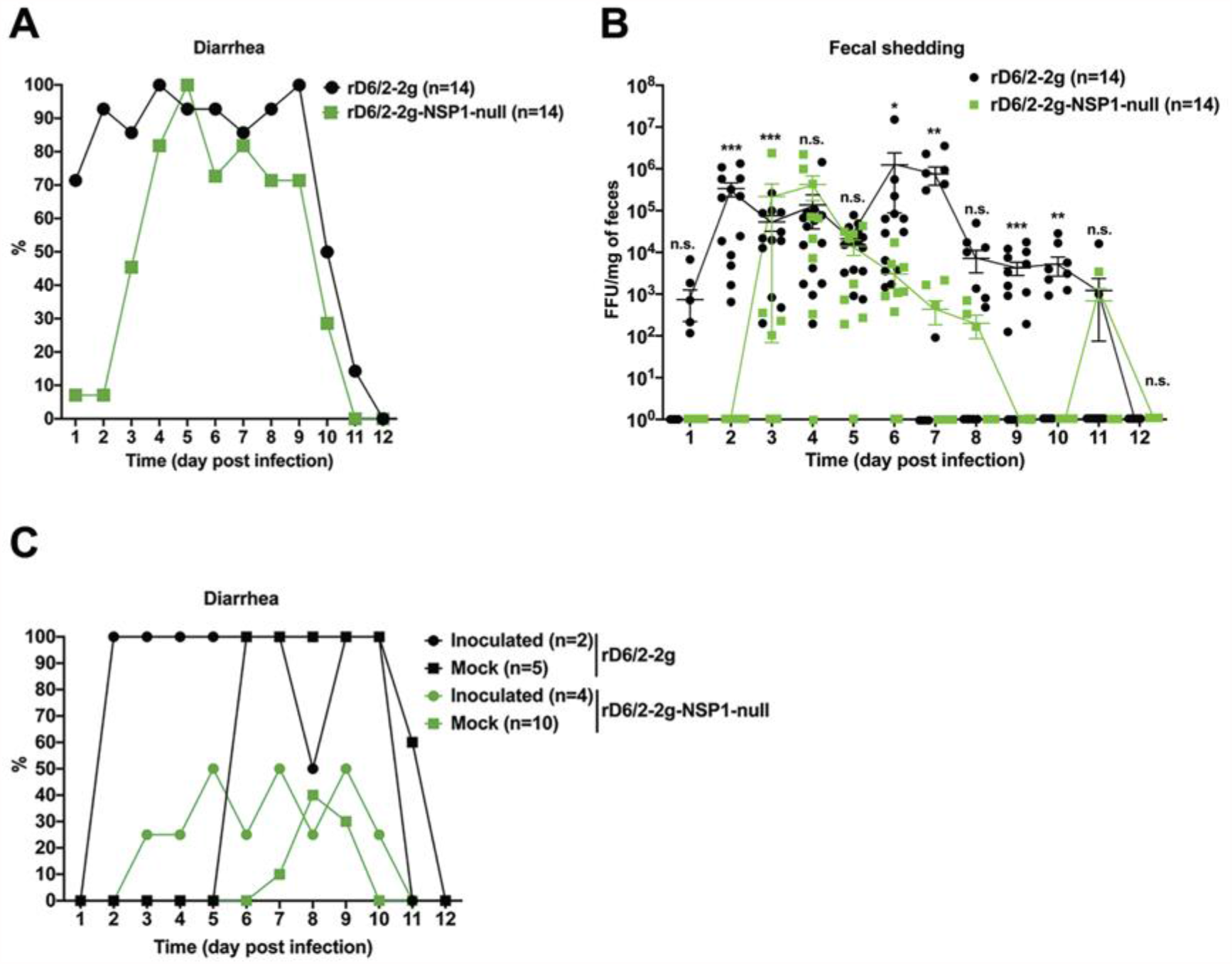
Characterization of diarrhea, fecal shedding, and transmission of rD6/2-2g and rD6/2-2g-NSP1-null in wild-type 129sv mice. (**A**) 5-day-old 129sv mice were orally inoculated with 1.5×10^3^ FFUs of rD6/2-2g and rD6/2-2g-NSP1-null. Diarrheal development was recorded from day 1 to 12 post infection. N indicates the number of mice used in each group. (**B**) Fecal shedding of the infectious RVs was monitored by an FFU assay and normalized by the weight of feces. Virus shedding within the same group on each day is shown as mean ± SEM. (**C**) To evaluate the transmission ability of the rD6/2-2g, 2 pups were orally infected with 1.5×10^3^ FFUs of rD6/2-2g, and 5 uninfected suckling littermates were co-housed with the inoculated pups in the same cage. For rD6/2-2g-NSP1-null, 2 pups in one cage and 2 in another cage were orally infected with the rD6/2-2g-NSP1-null described above, and co-housed with 6 and 4 uninoculated pups, respectively. Diarrhea was evaluated until 12 dpi as described above. * *P*<0.05; ** *P*<0.01; *** *P*<0.001; n.s., not significant (two-way ANOVA test).

To directly investigate whether NSP1 contributes to RV intestinal replication, we collected all three small intestinal segments (i.e., duodenum, jejunum, and ileum) from rD6/2-2g and rD6/2-2g-NSP1-null infected pups at 2 dpi and measured viral loads by RT-qPCR. The number of rD6/2-2g genome copies in the ileum was significantly (about 3 logs) higher than that of rD6/2-2g-NSP1-null, despite no major differences in the duodenum and jejunum (**Fig. 5A**). We also found that the viral loads of rD6/2-2g in mesenteric lymph nodes (MLNs) were higher than that of rD6/2-2g-NSP1-null, whereas no difference were observed in the blood, bile ducts, or the liver (**Fig. 5B-E**).

**Figure 5.**
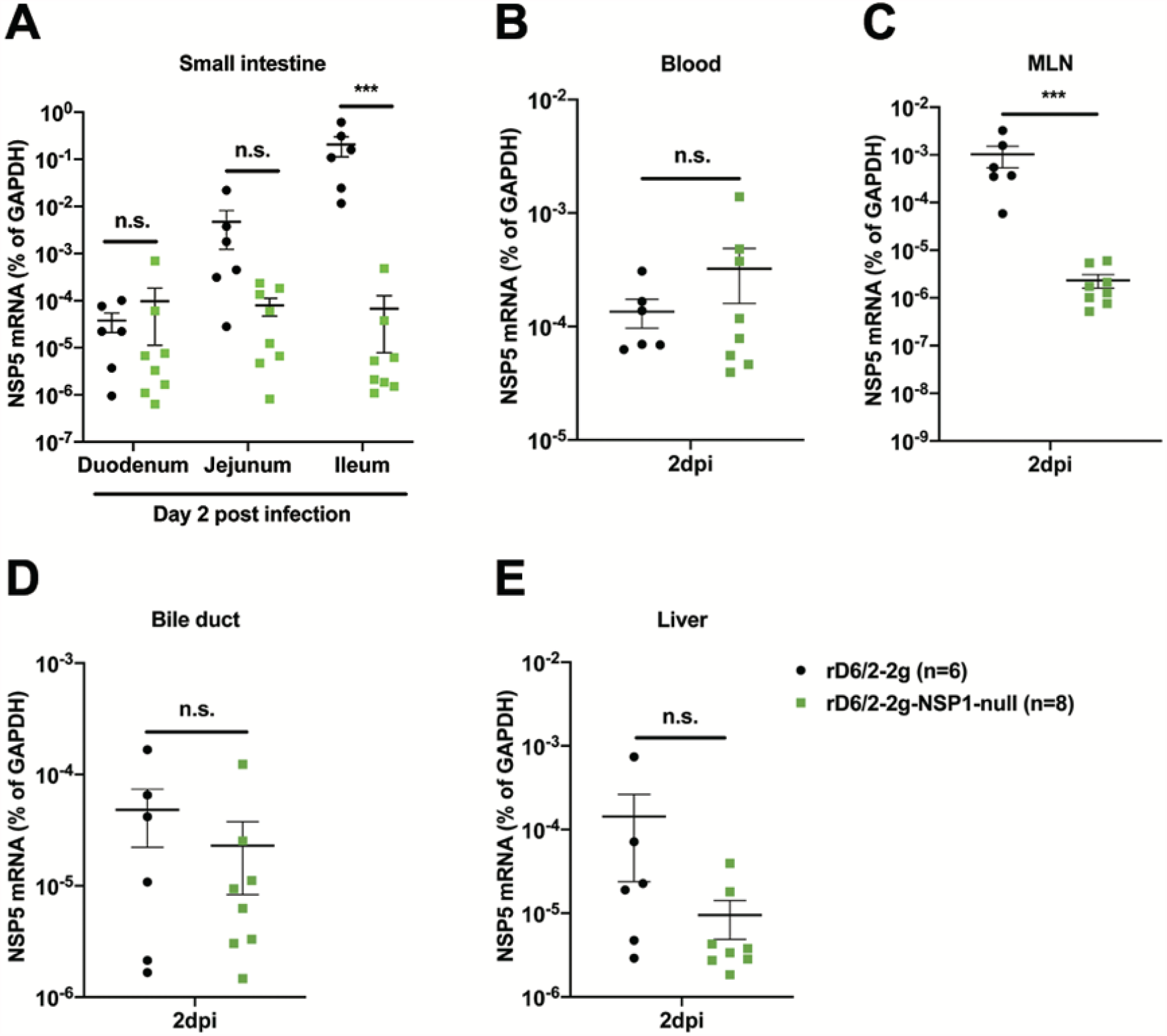
Viral loads of rD6/2-2g and rD6/2-2g-NSP1-null in the small intestines and indicated systemic sites in wild-type 129sv mice. (**A**) 5-day-old wild-type 129sv pups were orally infected with 1.5×10^3^ FFUs of rD6/2-2g and rD6/2-2g-NSP1-null. RNA was extracted from duodenum, jejunum, and ileum collected at 2 dpi and RT-qPCR was used to detect RV NSP5 mRNA levels. (**B-E**) Same as (**A**) except that blood, MLNs, bile duct, and liver were collected instead. *** *P*<0.001; n.s., not significant (unpaired student’s *t* test).

### Revertant NSP1 mutations rescued NSP1-deficient murine RV replication in mice

Notably, we noticed that 4 out of 14 mice from the rD6/2-2g-NSP1-null infected group shed high amount of infectious viruses in the feces at 3 dpi (**Fig. 4B**). This was reproducibly observed in two independent sets of experiments (6 out of 7 at 4 dpi in one experiment and 4 out of 7 at 3 dpi in the other experiment) (**Fig. S1**). To account for this enhanced replication, we amplified the 5’ end of gene 5 (17 to 466 nucleotides) directly from the stool samples collected from mice that had detectable RV shedding from 1-4 dpi and performed Sanger sequencing. Of interest, NSP1 fragments amplified from the rD6/2-2g-NSP1-null infected mice were indistinguishable from those from rD6/2-2g infection (**Table 1 and Fig. S2**), indicating that RV in the fecal specimens readily reverted back to the wild-type sequences. While we could not detect any RV shedding from rD6/2-2g-NSP1-null infected mice at 1 or 2 dpi, 3 out of the 4 mice that shed infectious RVs at 3 dpi had complete NSP1 reversion mutations and 1 had incomplete reversion (**Table 1 and Fig. S2**). 6 out of 14 mice infected with rD6/2-2g-NSP1-null shed virus at 4 dpi and all 6 animals had wild-type NSP1 sequences (**Table 1 and Fig. S2**). These data are consistent with our observation that the rD6/2-2g-NSP1-null infected mice unexpectedly developed diarrhea starting from 3 dpi and further emphasize an indispensable role that NSP1 protein plays during virus replication *in vivo*.

**Table 1.** NSP1 sequencing results from the fecal samples of wild-type and *Stat1* KO mice infected with rD6/2-2g-NSP1-null. Complete, 100% of both stop codons were completely reverted to wild-type sequences; mixed, only a portion of the viruses carrying the stop codons reverted to wild-type sequences.

| NSP1 Reversion Ratio |  |  |  |  |  |
| --- | --- | --- | --- | --- | --- |
|  | Reversion | Day 1 | Day 2 | Day 3 | Day 4 |
| **WT 129sv** | Complete |  |  | 3/4 | 6/6 |
|  | Mixed |  |  | 1/4 | 0/6 |
| **Stat1 KO**<br>**129sv** | Complete | 2/8 | 2/7 | 8/9 | 8/8 |
|  | Mixed | 6/8 | 5/7 | 1/9 | 0/8 |

### The blunted replication and pathogenesis of the recombinant NSP1-deficient murine RV is only partially recovered in the *Stat1* knockout mice

Since NSP1 functions as a highly potent IFN antagonist *in vitro*, we reasoned that rD6/2-2g-NSP1-null may be attenuated in wild-type 129sv mice due to the lack of IFN inhibitory capacity. To test this hypothesis, we orally infected 5-day-old *Stat1* knockout (KO) 129sv suckling pups, unable to respond to type I, II, and III IFNs, with 1.5×10^3^ FFUs of rD6/2-2g or rD6/2-2g-NSP1-null. For both viruses, the overall diarrheal development patterns in infected *Stat1* KO 129sv mice resembled those in wild-type mice (**Fig. 6A**). However, compared to rD6/2-2g infected animals that developed 57% and 100% diarrhea on 1 and 2 dpi, respectively, there was little to no diarrhea from rD6/2-2g-NSP1-null inoculated animals (**Fig. 6A**). We also evaluated the fecal RV shedding in the infected mice by an FFU assay. Even in the absence of IFN signaling, at 1 and 2 dpi, RV shedding of rD6/2-2g remained > 3 logs higher than that from the rD6/2-2g-NSP1-null infection (**Fig. 6B**), suggesting that NSP1 may facilitate RV replication in an IFN-independent manner.

**Figure 6.**
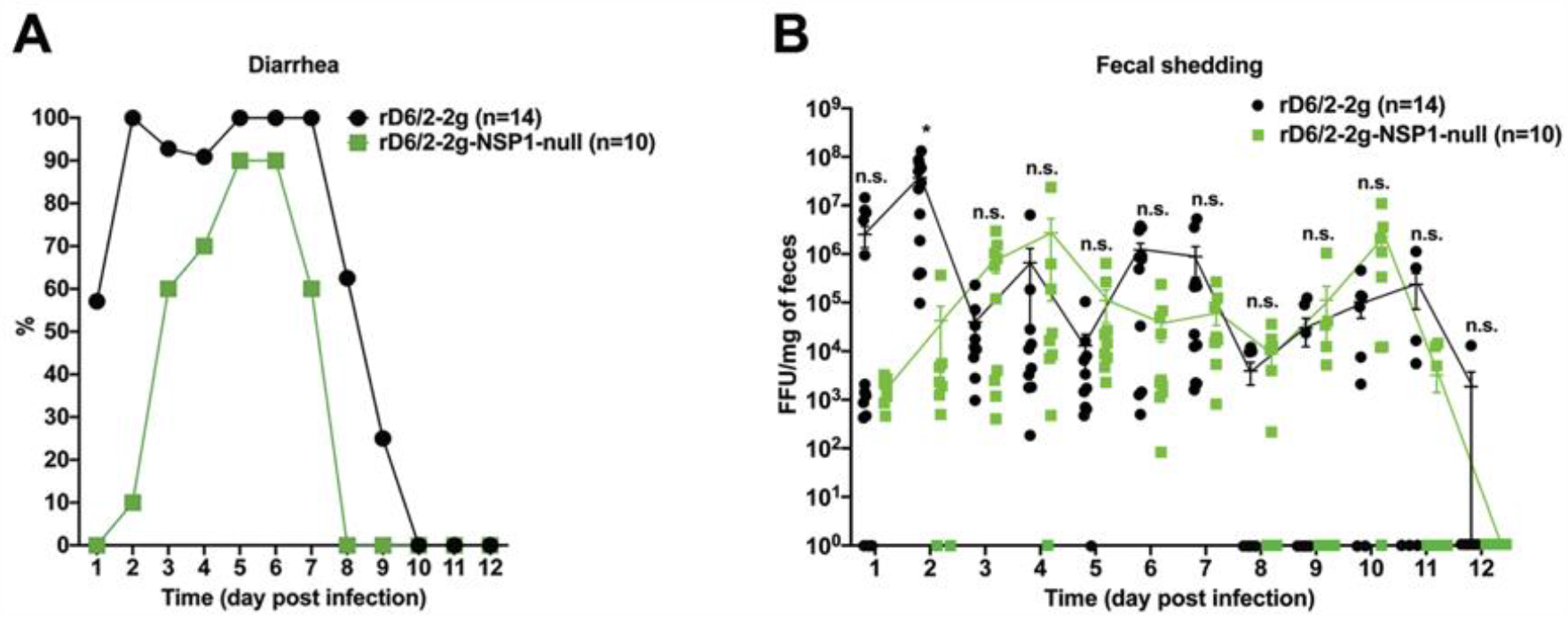
Characterization of diarrhea and fecal shedding of rD6/2-2g and rD6/2-2g-NSP1-null in *Stat1* KO 129sv mice. (**A**) 5-day-old *Stat1* KO 129sv mice were orally inoculated with 1.5×10^3^ FFUs of rD6/2-2g and rD6/2-2g-NSP1-null. The diarrhea rate was monitored from day 1 to 12 post infection. (**B**) Viral shedding in stool samples were detected by an FFU assay, and normalized by the feces weight. Virus shedding within one group on each day is shown as mean ± SEM. * *P*<0.05; n.s., not significant (two-way ANOVA test).

Consistent with the diarrhea and fecal shedding results, we found that for all the small intestinal tissues examined, rD6/2-2g-NSP1-null was still severely attenuated compared to rD6/2-2g in *Stat1* KO mice at 2 dpi (**Fig. 7A**). The viral loads of rD6/2-2g in the ileum were approximately 7,000-fold higher than those of rD6/2-2g-NSP1-null infected mice (**Fig. 7A**). Furthermore, rD6/2-2g also had significantly more spread to systemic organs including the blood, MLNs, bile duct, and liver than rD6/2-2g-NSP1-null (**Fig. 7B-E**). Collectively, these data indicate that even in a host devoid of IFN signaling, the replication of a murine RV without NSP1 expression remains highly attenuated *in vivo*.

**Figure 7.**
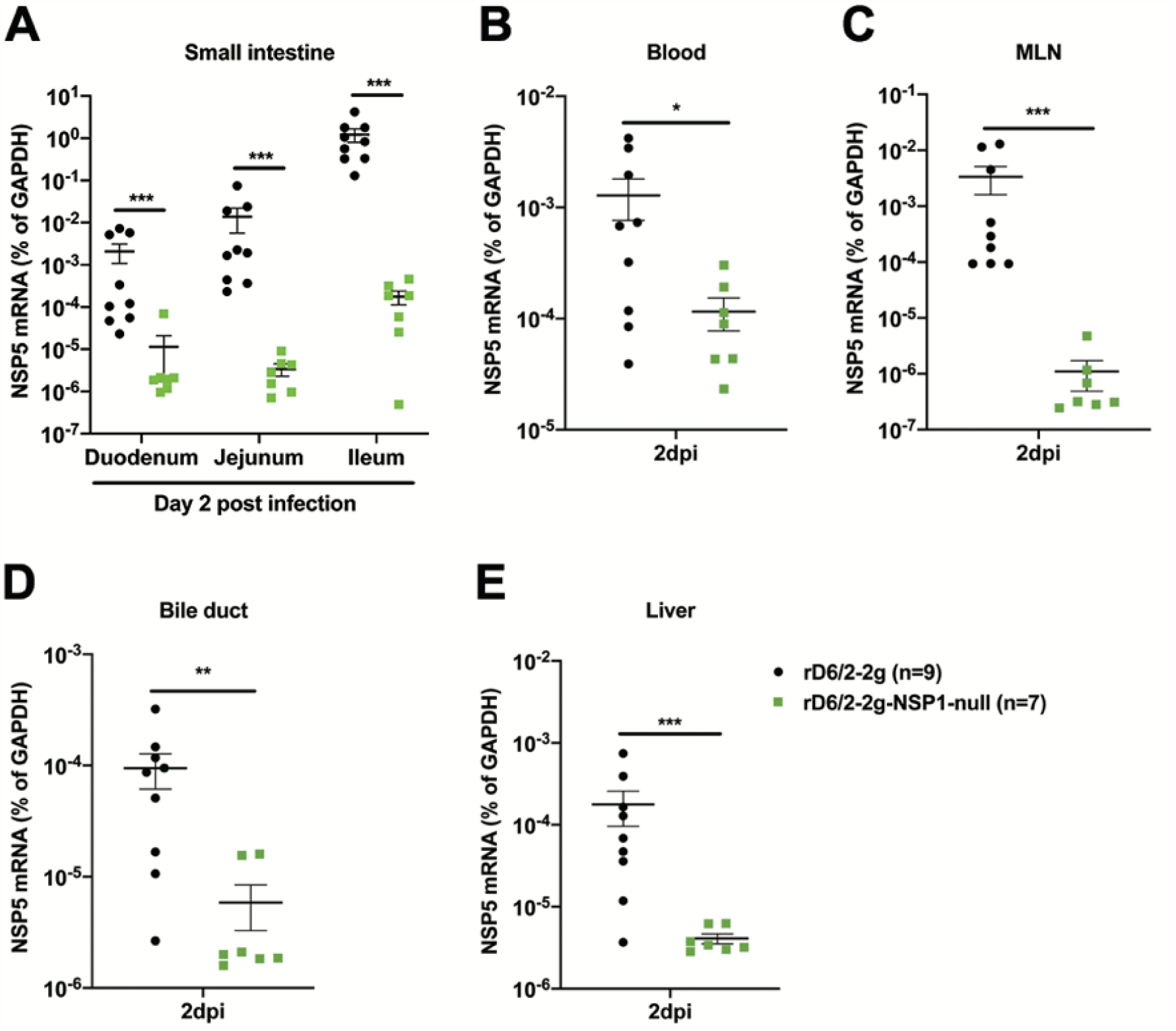
Viral loads of rD6/2-2g and rD6/2-2g-NSP1-null in the small intestines and indicated extra-intestinal tissues in *Stat1* KO 129sv mice. (**A**) 5-day-old *Stat1* KO 129sv pups were orally infected with 1.5×10^3^ FFUs of rD6/2-2g and rD6/2-2g-NSP1-null. The duodenum, jejunum, and ileum were collected at 2 dpi and RV NSP5 mRNA levels were detected by RT-qPCR. (**B-E**) Same as (**A**) except that blood, MLNs, bile duct, and liver were collected instead. * *P*<0.05; ** *P*<0.01; *** *P*<0.001; n.s., not significant (unpaired student’s *t* test).

Further supporting an IFN-independent role of NSP1 is the fact that rD6/2-2g-NSP1-null also reverted back to the wild-type sequences in *Stat1* KO mice. We surveyed over 30 fecal samples from the rD6/2-2g-NSP1-null infected mice. Out of the 8 mice that shed infectious RVs at 1 dpi, 2 had complete NSP1 revertant mutations and the other 6 mice had incomplete reversion (**Table 1 and Fig. S2**). At 4 dpi, 8 out of the 10 mice that have substantial fecal shedding had viruses with wild-type NSP1 sequences (**Table 1 and Fig. S2**). Therefore, all NSP1-deficient viruses reached 100% reversion in *Stat1* KO 129sv mice by 4 dpi, the same as that in the wild-type 129sv mice.

### The attenuated replication of the recombinant NSP1-deficient simian RV SA11 is not rescued in *Stat1* KO mice

To further corroborate the unexpected finding that NSP1 is required for RV replication in *Stat1* KO mice, we turned to the heterologous simian RV SA11 strain that leads to low but detectable fecal shedding when the mice are inoculated at a high dose (1×10^7^ PFUs) (21). We successfully rescued an NSP1-deficient SA11 (rSA11-NSP1-null), which was given to 5-day-old wild-type and *Stat1* KO 129sv mice in direct comparison to the parental rSA11 at 1×10^7^ PFUs. The overall trends of diarrheal development between rSA11 and rSA11-NSP1-null in wild-type 129sv mice were similar (**Fig. 8A**). In *Stat1* KO mice, both viruses developed less diarrhea than in the wild-type animals but the rSA11-NSP1-null seemed to be even more attenuated than rSA11 (**Fig. 8A**). We also quantified RV antigen shedding in the stool samples collected from rSA11 or rSA11-NSP1-null infected mice by an enzyme-linked immunosorbent assay (ELISA). While neither virus had detectable shedding in wild-type pups, we observed that rSA11 but not rSA11-NSP1-null resulted in transient shedding between 4-6 dpi (**Fig. 8B**). Consistently, we found higher viral loads in the small intestines of rSA11 infection compared to rSA11-NSP1-null infection of *Stat1* KO mice (**Fig. 8C**). Taken together, these data suggest that despite limited replication ability, the heterologous simian RV SA11 strain is also attenuated without NSP1 expression in IFN-deficient animals, resembling what we observed with homologous murine RV infections.

**Figure 8.**
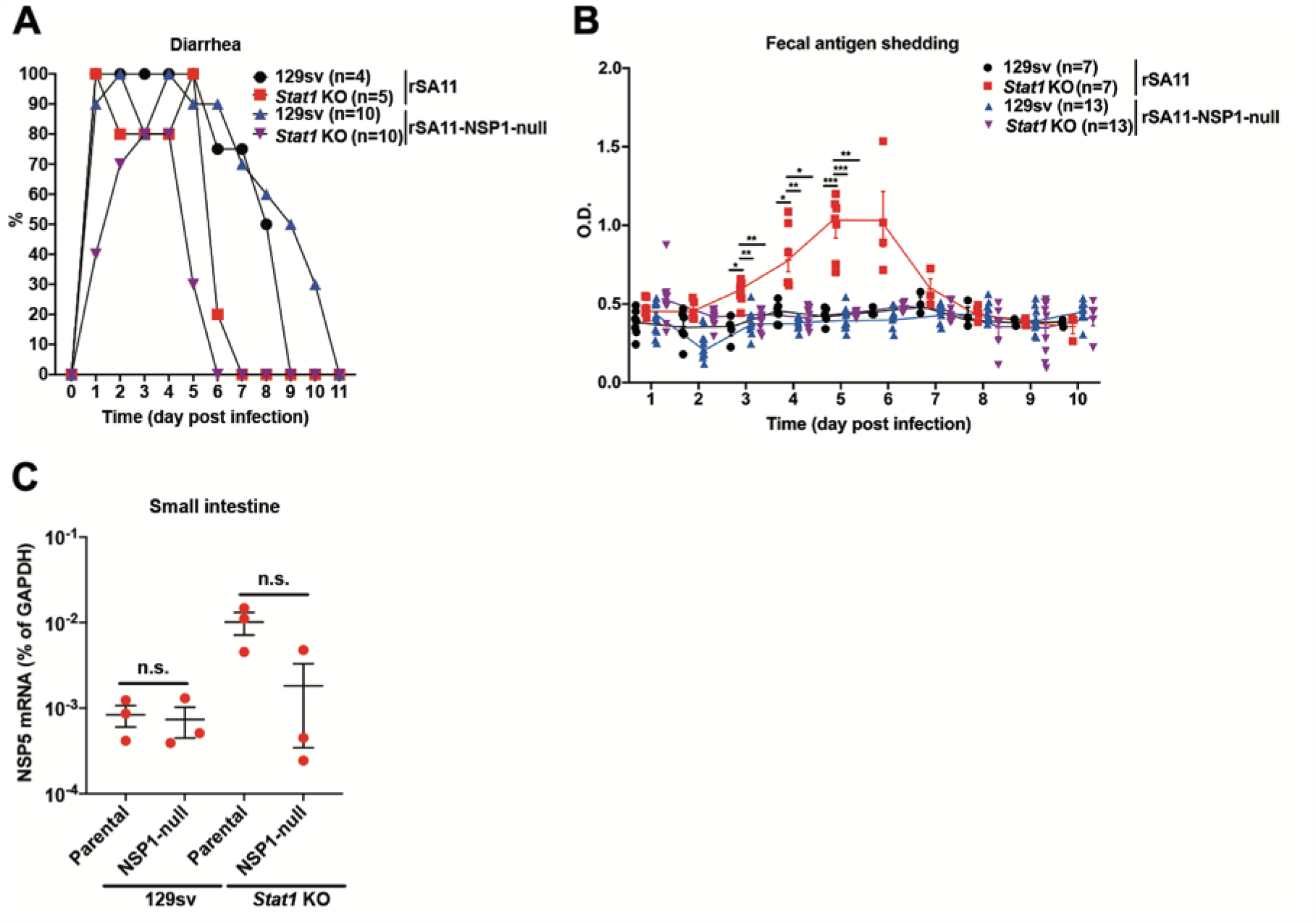
Characterization of diarrhea, fecal shedding, and replication of heterologous rSA11 and rSA11-NSP1-null in wild-type and *Stat1* KO 129sv mice. (**A**) 5-day-old wild-type and *Stat1* KO 129sv mice were orally infected with 1×10^7^ PFUs of rSA11 and rSA11-NSP1-null. Diarrheal development was recorded from day 0 to 11 post inoculation. (**B**) Stool samples were collected from 1 to 12 days post infection, and virus shedding in feces was measured by ELISA and plotted as optical density (O.D.) values. * *P*<0.05; ** *P*<0.01; *** *P*<0.001; n.s., not significant (two-way ANOVA test). (**C**) Small intestinal tissues were collected at 5 dpi. Total RNA was extracted and RV NSP5 mRNA levels were detected by RT-qPCR. n.s., not significant (unpaired student’s *t* test).

## DISCUSSION

In this study, we exploited a recently developed and further optimized plasmid-based RG system to examine the role of NSP1 in RV replication *in vivo*. Previous studies showed that naturally isolated RV SA11-5S and SA11-30-1A variants (24), and some RG rescued recombinant RVs, albeit unable to induce IRF3 degradation, replicate efficiently in cell culture (20). However, because of the limited replication and transmission of non-murine heterologous RVs in mice (25), until now it is difficult to assess whether NSP1 is required for RV replication *in vivo*. Here, taking advantage of an optimized RG system, we rescued an NSP1-deficient murine like rD6/2-2g-NSP1-null (**Fig. 1**), which replicated similarly to rD6/2-2g in MA104 cells (**Fig. 2A and B**). The plaque size formed by rD6/2-2g-NSP1-null was, however, much smaller than that of rD6/2-2g (**Fig. 2C**), reminiscent of the data derived from the SA11 strain (19, 20). The smaller plaque size may reflect a defect in the efficiency of cell-cell spread.

The similar growth properties of rD6/2-2g-NSP1-null and rD6/2-2g in MA104 may be due to IFN-defective MA104 cells, which also supported comparable propagation of the parental rSA11 strain and recombinant rSA11-dC103 and rSA11-Nluc that both lack an intact NSP1 expression. The replication of rSA11-Nluc in IFN-competent cells was, however, much lower than that of rSA11 (19, 26), and this phenotype may be caused by the missing IRF3 degradation ability of rSA11-dC103 and rSA11-Nluc. Thus, in order to assess whether NSP1 was important to murine RV replication in IFN-competent cells, we examined the growth curves of rD6/2-2g-NSP1-null and rD6/2-2g in HEK293 and HAP1 cells. Intriguingly, these two viruses still exhibited similar replication properties (**Fig. 3**). Although we observed a 3-4-fold higher titer of rD6/2-2g in HAP1 cells at 48 h post infection, there were no statistically significant differences at any other time points (**Fig. 3**). These results raise the possibilities that another RV protein may compensate for the loss of NSP1 or that the IFN-mediated antiviral activities are both cell type and virus strain-specific.

Recent studies have reported that the recombinant murine-like RV (rD6/2-2g) replicate robustly in the small intestines of 129sv suckling pups, that rD6/2-2g infection causes diarrheal diseases, and that the transmission efficiency of rD6/2-2g is similar to that of the original reassortant murine RV D6/2 strain (22). Here, we orally inoculated the 129sv mice with rD6/2-2g or rD6/2-2g-NSP1-null. The lower percentage of diarrheal disease and lower titer of RV shedding in rD6/2-2g-NSP1-null infected pups at the early time points revealed that NSP1 protein is important for virus pathogenesis *in vivo*. However, starting on day 3 post infection, the curves of diarrhea and virus shedding of rD6/2-2g-NSP1-null approached those of rD6/2-2g parental strain. This observation led us to ask whether the rD6/2-2g-NSP1-null reverted to wild type rD6/2-2g, despite the presence of two stop codons inserted at the very beginning of NSP1 open reading frame. To our surprise, all of the gene 5 segments amplified from rD6/2-2g-NSP1-null infected 129sv mice examined at 4 dpi had completely reverted to wild-type sequences. We performed the infection experiments with rD6/2-2g and rD6/2-2g-NSP1-null at different time points, and these data were repeated in two independent experiments (**Fig. S1**), thus the reversion observed in rD6/2-2g-NSP1-null inoculated mice is unlikely due to the contamination by rD6/2-2g. We also amplified NSP1 from fecal specimens obtained on days 1-4 post infection from *Stat1* KO 129sv mice. 2 out of 8 mice had complete NSP1 reversion mutations and the remaining 6 had incomplete reversion as early as 1 dpi; meanwhile, all the 8 *Stat1* KO 129sv mice had complete NSP1 reversion mutations at 4 dpi (**Table 1 and Fig. S2**). These results emphasized the essential role of NSP1 protein during RV replication *in vivo*. We found significantly higher levels of rD6/2-2g in the small intestines, blood, MLNs, bile duct, and liver than rD6/2-2g-NSP1-null infected *Stat1* KO 129sv mice on 2 dpi (**Fig. 7**), although the only statistically significant differences examined in the wild-type 129sv mice on 2 dpi were the ileum and MLNs (**Fig. 5**). We found that the replication of rD6/2-2g in small intestines and systemic organs in *Stat1* KO 129sv mice was about 10-fold higher than in wild-type 129sv mice, but the replication of rD6/2-2g-NSP1-null in these tissues in wild-type 129sv mice was similar to that of *Stat1* KO 129sv mice (**Fig. 5 and 7**). These data suggest that NSP1 protein contributes to virus replication but this contribution appears to be independent of IFN signaling during RV infection *in vivo*. We also found that the replication of the heterologous rSA11-NSP1-null and rSA11 simian RVs was comparable in wild-type 129sv mice, but the replication of rSA11 was increased in *Stat1* KO 129sv mice (**Fig. 8**). Unlike rD6/2-2g, we did not see a recovery of rSA11-NSP1-null in wild-type mice, probably because the replication level was too low to permit generation and selection of sufficient mutations by the viral polymerase.

In conclusion, this study identifies the importance of the NSP1 protein in promoting virus replication *in vivo*. Our data suggest that NSP1 not only inhibits the IFN response but may also acts to block some other antiviral signaling pathways or facilitates virus replication independent of IFN signaling. For instance, Nlrp9b inflammasome also restricts RV infection in intestinal epithelial cells, but whether this phenotype was related to NSP1 is still unknown (27). In future studies, it will be interesting to investigate the exact domains of NSP1 responsible for its pro-viral replication functions *in vivo*. In addition, since NSP1 is involved in RV host range restriction, one can study the NSP1 functionality in the context of a homologous virus infection by replacing the murine RV NSP1 with NSP1s derived from heterologous RV strains. With an optimized RG system and fully virulent murine RVs, we expect to uncover the physiological functions of other RV-encoded viral factors. We anticipate that a deeper understanding of the rotavirus-host interactions and rotavirus pathogenesis *in vivo* will guide the development of novel next-generation RV vaccine candidates with improved efficacy in developing countries and higher compatibility with an immunocompromised population.

## FIGURES and FIGURE LEGENDS

**Supplemental Figure 1.**
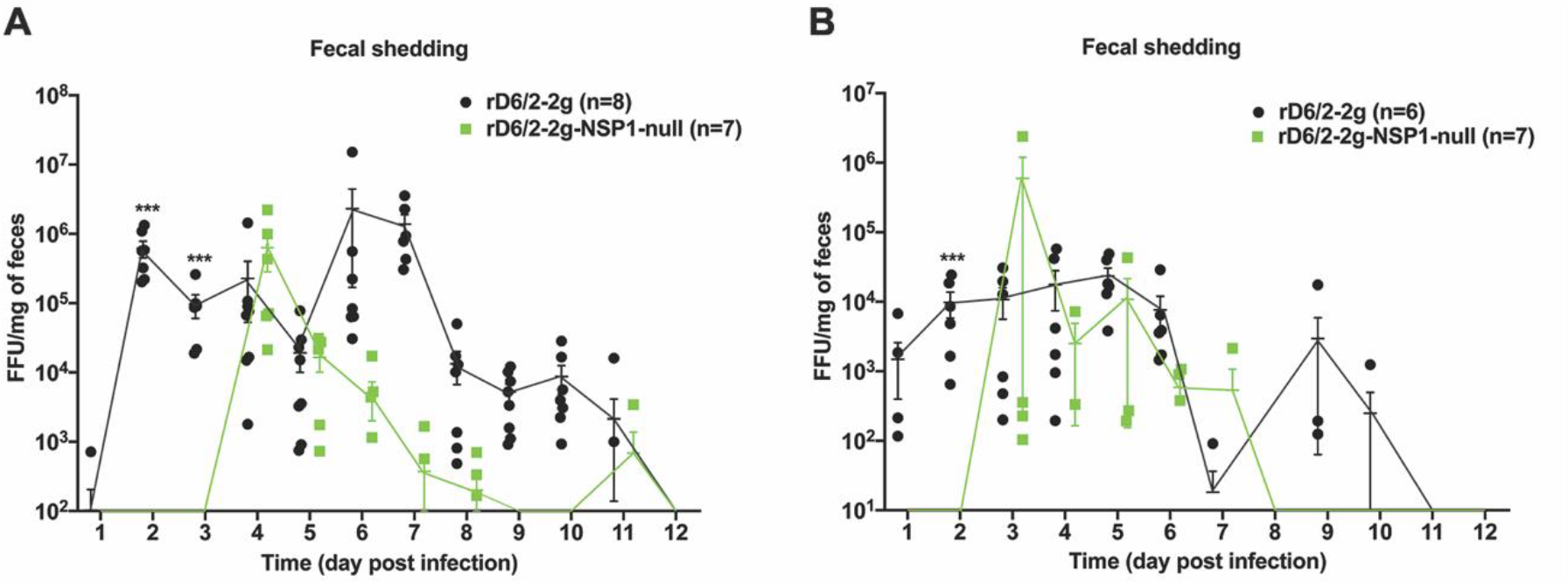
Characterization of the virus shedding of rD6/2-2g and rD6/2-2g-NSP1-null in wild-type 129sv mice. (**A**) 5-day-old 129sv mice were orally inoculated with 1.5×10^3^ FFUs of rD6/2-2g and rD6/2-2g-NSP1-null. Viral shedding in stool samples were detected by an FFU assay and normalized by the feces weight. (**B**) Another two cages of suckling mice were orally infected as in (**A**), and the fecal shedding was measured and normalized as described above. Virus shedding within the same group on each day is shown as mean ± SEM. *** *P*<0.001 (two-way ANOVA test).

**Supplemental Figure 2.**
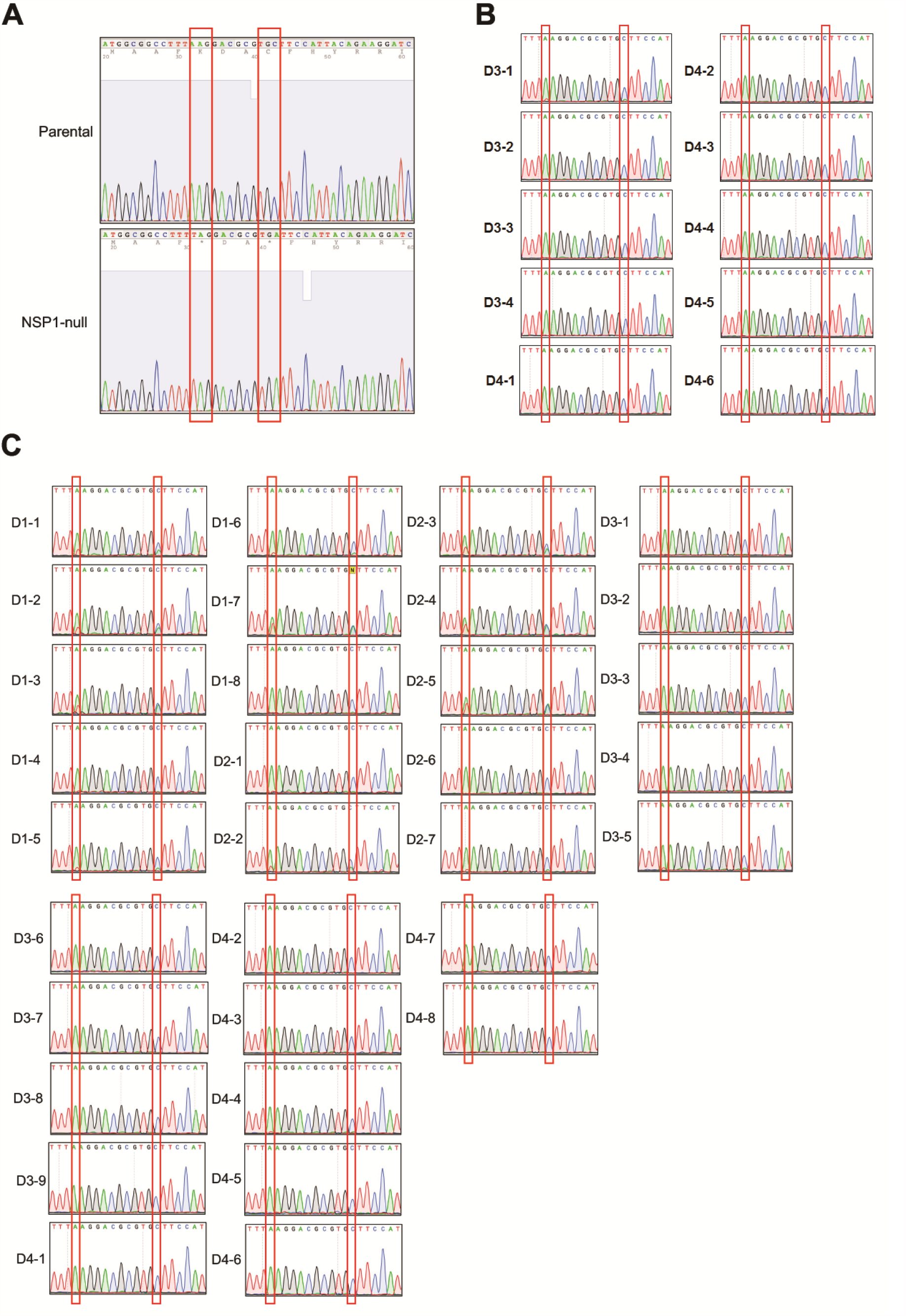
Sanger sequencing of RV gene 5 fragment. (**A**) NSP1 fragments (17 to 466 nucleotides) were amplified from the rD6/2-2g and rD6/2-2g-NSP1-null virus stocks and analyzed by Sanger sequencing. (**B**) Same as (**A**) except that feces from rD6/2-2g-NSP1-null inoculated wild-type 129sv mice at day 3 and 4 post infection were analyzed instead. (**C**) Same as (**A**) except that feces from rD6/2-2g-NSP1-null inoculated *Stat1* KO 129sv mice at day 1 to 4 post infection were analyzed instead.

## MATERIALS AND METHODS

### Cells and viruses

The rhesus monkey kidney epithelial MA104 cells were grown in medium 199 (Gibco) supplemented with 10% heat-inactivated fetal bovine serum (FBS) (VWR), 100 U/ml penicillin, 100 μg/ml streptomycin, and 0.292 mg/ml L-glutamine. BHK-T7 cell, a baby hamster kidney cell line stably expressing T7 RNA polymerase was kindly gifted by Ursula Buchholz (Laboratory of Infectious Diseases, NIAID, NIH, USA), and was cultured in Dulbecco’s modified Eagle’s medium (DMEM) (Gibco) supplemented with 10% heat-inactivated FBS, 100 U/ml penicillin, 100 μg/ml streptomycin, and 0.292 mg/ml L-glutamine, and also 0.3 mg/ml G418 (Promega) was added to the culture medium at every other passage. MA104 cells stably expressing parainfluenza virus 5 V protein and bovine viral diarrhea virus N protease were cultured in medium 199 supplemented with 10% heat-inactivated FBS, 100 U/ml penicillin, 100 μg/ml streptomycin, 0.292 mg/ml L-glutamine. 10 μg/ml blasticidin and 10 μg/ml puromycin were added to the medium at every other passage as described previously (22).

The simian RV SA11 strain and the murine RV D6/2 strain were propagated as described previously (7, 19). Briefly, RV stock was activated with 5 μg/ml trypsin (Gibco Life Technologies, Carlsbad, CA) for 20 min at 37°C. The activated RV were incubated with MA104 cells, which were washed twice with serum free medium 199 for 1 h at 37°C. Then the viruses were removed and the new serum free medium 199 supplemented with 0.5 μg/ml trypsin were added to the MA104 cells. Virus titers were determined by a standard plaque assay in MA104 cells.

### Sequencing of the murine RV D6/2 strain

Viral particle enrichment of cell culture supernatant was performed based on the NetoVIR protocol (28). Briefly, cell culture supernatant was centrifuged for 3 min at 17,000 × *g* and filtered with a 0.8-μm PES filter (Sartorius). The filtrate was treated with benzonase (Novagen) and micrococcal nuclease (New England BioLabs) at 37°C for 2 h to remove the free-floating nucleic acids. Subsequently, samples were extracted using the QIAamp Viral RNA minikit (Qiagen) according to the manufacturer’s instructions, without addition of carrier RNA to the lysis buffer. Reverse transcription and second strand synthesis were performed by an adjusted version of the whole-transcriptome amplification (WTA2) protocol (Sigma-Aldrich), as described previously (29). A sequencing library was constructed with the Nextera XT library preparation kit (Illumina). The size of the library was checked with Bioanalyzer (Agilent Technologies) with a high sensitivity DNA chip and the 2 nM pooled libraries were sequenced on an Illumina NextSeq 500 platform (2 × 150 bp paired ends).

Low-quality reads, ambiguous bases, and primer and adapter sequences were removed from the paired-end reads with Trimmomatic with default parameters (30). Trimmed reads were *de novo* assembled with metaSPAdes from SPAdes software using 21, 33, 55, and 77 k-mer lengths (31). The obtained contigs were annotated with DIAMOND against a nonredundant protein database (32). Contigs annotated as “rotavirus” were extracted. The obtained sequences were verified *in silico* by remapping the trimmed reads to the obtained contigs using BWA software (33).

### Construction of a T7 plasmid encoding mutant gene 5 of D6/2 or SA11

To rescue an NSP1-deficient murine-like RV, we generated a pT7-D6/2-NSP1-null via the QuikChange II site-directed mutagenesis kit (Agilent Technology) based on pT7-D6/2-NSP1 (22). Briefly, the AAG and TGC codons in the NSP1 ORF of D6/2 were replaced with stop codons TAG and TGA at nucleotide position 43 to 45, and 52 to 55 in pT7-D6/2-NSP1. The mutant primers used were: rD6/2-2g-NSP1-null forward primer: 5’-GTGTTAGCCATGGCGGCCTTTTAGGACGCGTGATTCCATTACAGAAGG-3’; rD6/2-2g-NSP1-null reverse primer: 5’-CCTTCTGTAATGGAATCACGCGTCCTAAAAGGCCGCCAT GGCTAACAC-3’. To rescue the NSP1-deficient simian RV SA11, we generated the plasmid pT7-SA11-NSP1-null using the same strategies described above. The mutant primers used were: SA11-NSP1-null forward primer: 5’-GCTACTTTTAAAGATGCATGCTTTTAATAGCGTAGATTAACTGCTTTAAATCGG-3’; SA11-NSP1-null reverse primer: 5’-CCGATTTAAAGCAGTTAATCTACGCTATTAAAAGCATGCATCTTTAAAAGTAGC-3’.

### Generation of recombinant murine like and simian RVs

Recombinant rD6/2-2g and rD6/2-2g-NSP1-null were generated according to the optimized entirely plasmid-based RG system described recently (22). Briefly, 0.4 μg of pT7-SA11-VP1, pT7-D6/2-VP2, pT7-D6/2-VP3, pT7-D6/2-VP4, pT7-D6/2-VP6, pT7-D6/2-VP7, pT7-D6/2-NSP1 (or pT7-rD6/2-NSP1-null), pT7-D6/2-NSP3, and pT7-SA11-NSP4, 1.2 μg of pT7-D6/2-NSP2 and pT7-D6/2-NSP5, 0.8 μg of the helper plasmid C3P3-G1, and 14 μl TransIT-LT1 (Mirus) transfection reagent were mixed together and transfected into BHK-T7 cells in 12-well plate. 18 h later, the transfected BHK-T7 cells were washed twice with FBS-free DMEM, then supplemented with 800 μl fresh FBS-free DMEM, 24 h later, 1×10^5^ MA104 N*V cells in 200 μl FBS-free DMEM along with 0.5 μg/ml trypsin were added to the transfected BHK-T7 cells for another 3 days. After that, mixed cells were frozen and thawed for 3 times. The rescued virus was propagated for two passages in MA104 cells in 6-well plate, then the virus was propagated in T75 flask to produce the virus stock.

The recombinant SA11 and SA11-NSP1-null were rescued using the same protocol described above, expect that the plasmids used were pT7-SA11-VP1, pT7-SA11-VP2, pT7-SA11-VP3, pT7-SA11-VP4, pT7-SA11-VP6, pT7-SA11-VP7, pT7-SA11-NSP1 (or pT7-SA11-NSP1-null), pT7-SA11-NSP2, pT7-SA11-NSP3, pT7-SA11-NSP4, and pT-SA11-NSP5.

### Purification of RV particles by sucrose gradient centrifugation

RVs were concentrated by sucrose cushion as described previously (34). Briefly, RVs propagated in T75 flasks were harvested by repeat the freeze-thaw cycle three times. Then we clarified the crude lysate of cell debris by centrifugation at 3,000 × g for 1 h at 4°C. After that, 8 ml of clarified RVs were placed into a 10 ml SW44 ultracentrifuge tube, and 2 ml 40% sucrose (w/v) was carefully added to the bottom of the ultracentrifuge tube, and subjected to centrifugation at 35,000 rpm for 3 h at 4°C. At last, we removed the supernatant, and add 200 μl FBS-free medium 199 to resuspend the concentrated RVs at 4°C overnight.

### Electrophoretic analysis of viral genomic dsRNAs

Viral genomic dsRNAs were extracted from sucrose cushion concentrated viruses using TRIzol reagent (Thermo Scientific) according to the manufacture’s protocol (35). Then the dsRNAs were mixed with gel loading dye, purple (6×) (New England Biolabs). Samples were loaded into a 4-15% precast polyacrylamide gel, and running for 3 h at 180 volts. Gel was stained for 1 h with 0.1 μg/ml ethidium bromide and visualized by ChemiDoc MP Imaging System (Bio-Rad).

### Restriction enzyme digestion and sequencing analysis

The total RNA of the recombinant rD6/2-2g, rD6/2-2g-NSP1-null virus stocks, and stool samples was extracted by TRIzol. Total RNA was reverse transcribed to complementary DNA using High Capacity cDNA Reverse Transcription Kit with RNase Inhibitor (Applied Biosystems) according to the user guide. Briefly, 0.8 μg of RNA, 2 μl of 10× RT Buffer, 0.8 μl of 100 mM dNTP Mix, 2 μl of RT random primers, 0.1 μl of RNase Inhibitor, 0.1 μl of MultiScribe Reverse Transcriptase, and flexible amount of nuclease-free H_2_O were added to the 20 μl reaction. Reverse transcription thermocycling program was set at 25°C for 10 min, 37°C for 2 h, and 85°C for 5 min.

NSP1 5’ end fragments were amplified by Phusion Hot Start II DNA Polymerase (Thermo Scientific) following the manufacturer’s guide. The primers used for PCR were: NSP1 forward primer: 5’-GTCTTGTGTTAGCCATGGC-3’, NSP1 reverse primer: 5’-CAGCGGTTAAAGTGATCGG-3’. PCR products were gel-purified using QIAquick Gel Extraction Kit (QIAGEN). 0.1 μg of NSP1 fragments were digested by restriction enzyme *HinfI* (NEB) for 1 h at 37°C. The enzyme digested products were separated by 2% agarose gel electrophoresis, stained by ethidium bromide, and visualized by ChemiDoc MP Imaging System (Bio-Rad). A separate set of purified NSP1 fragments were sent for Sanger sequencing using the NSP1 reverse primer.

### Immunoblotting

MA104 cells in 24-well plate were infected by rD6/2-2g or rD6/2-2g-NSP1-null at an MOI of 3 for 6 h. Then uninfected and infected MA104 cells were washed twice by ice-cold phosphate-buffered saline (PBS; Thermo Scientific), and lysed in RIPA buffer (150 mM NaCl, 1.0% IGEPAL CA-630, 0.5% sodium deoxycholate, 0.1% SDS, 50 mM Tris, pH 8.0; Sigma-Aldrich) supplemented with 1× protease inhibitor cocktail (Thermo Scientific) for 30 min at 4°C. After that, cell debris was removed by centrifuge at 12,000 ×g for 10 min at 4°C. Samples were resolved in precast SDS-PAGE gel (4-15%; Bio-Rad) and transferred to nitrocellulose membrane (0.45 μm; Bio-Rad). The membrane was incubated with blocking buffer (5% bovine serum albumin (BSA) diluted in PBS supplemented with 0.1% Tween 20) for 1 h at room temperature. Then the membrane was incubated with anti-IRF3 rabbit monoclonal antibody (CST, #4302, 1:1000), anti-RV VP6 mouse monoclonal antibody (Santa Cruz Biotechnology, sc-101363, 1:1000), anti-GAPDH rabbit monoclonal antibody (CST, #2118, 1;1000), followed by incubation with anti-mouse IgG (CST, #7076, 1:5000) or anti-rabbit IgG (CST, #7074, 1:5000) horseradish peroxidase-linked (HRP) antibodies. The antigen-antibody complex was detected using Clarity Western ECL substrate (Bio-Rad), and ChemiDoc MP Imaging System according to the manufacturer’s manuals.

### RT-QPCR

RT-qPCR was performed using the above cDNA as described previously (23). The expression level of housekeeping gene GAPDH was quantified by 2× SYBR Green Master Mix (Applied Biosystems), and NSP5 was measured by 2× TaqMan Fast Advanced Master Mix (Applied Biosystems). The primers used in this study were as follows: human GAPDH forward primer: 5’-GGAGCGAGATCCCTCCAAAAT-3’, reverse primer: 5’-GGCTGTTGTCATACTTCTCATGG-3’; mouse GAPDH forward primer: 5’-TCTGGAAAGCTGTGCCGTG-3’, reverse primer: 5’-CCAGTGAGC TTCCCGTTCA G-3’; NSP5 forward primer: 5’-CTGCTTCAAACGATCCACTCAC-3’, reverse primer: 5’-TGAATCCATAGACACGCC-3’, probe: 5’-CY5/TCAAATGCAGTTAAGACAAATGCAGACGCT/IABRQSP-3’.

### Plaque assay

Plaque assay was performed as described previously (36). Briefly, 1×10^5^ cells/ml MA104 cells were seeded in 6-well plate and virus samples were serially diluted 10-fold and incubated with the confluent MA104 cells for 1 h at 37°C. Then samples were replaced by FBS-free medium 199 with 0.1% agarose supplemented with 0.5 μg/ml trypsin and put back to 37°C. Plaques were visualized at day 3 to day 5 post inoculation by 0.0165% neutral red staining. In order to measure the size of the plaques, we recorded more than 75 plaques by the microscope (ECHO) in two different experiments. Then the diameters of the plaques were calculated by the annotation tool of the microscope.

### Focus-forming unit assay

Focus-forming assay was conducted as described previously (35). Briefly, 1×10^5^ cells/ml MA104 cells were seeded in 96-well plate, then virus samples were serially diluted 5 or 10-fold and incubated with a monolayer of MA104 cells for 10 h or overnight at 37°C. Then cells were fixed by 10% formalin, followed by permeabilized with 1% Tween 20. After that cells were incubated with anti-rotavirus capsid mouse monoclonal antibody and anti-mouse HRP-linked antibodies. The foci were stained by 3-amino-9-ethylcarbazole HRP substrate (Vector laboratories) and stopped by wash twice with PBS.

### Mice infection

Wild-type 129S1/SvImJ or *Stat1* KO mice were purchased from the Jackson Laboratory and Taconic Biosciences, respectively, and bred locally at the WUSTL BJCIH vivarium. 5-day-old suckling pups were orally infected with rescued rD6/2-2g (1.5×10^3^ FFU) and rD6/2-2g-NSP1-null (1.5×10^3^ FFU) (37). Diarrhea was evaluated from day 1 to day 12 post infection. In the meantime, feces from infected mice were also collected, and focus-forming assay was used to titrate RV in stool samples. Briefly, 50 μl PBS with calcium and magnesium was added to the 1.5 ml Eppendorf tubes, and the weight was recorded. After we collected the feces, these tubes were weighted again, and the stool samples were homogenized before we made the serial dilution to conduct the focus-forming assay. Duodenum, jejunum, ileum, blood, mesenteric lymph node, and liver were collected from inoculated pups at day 2 post infection, and immediately placed in liquid nitrogen and stored at -80°C until use. RNA was extracted from those tissues using RNeasy Plus Mini Kit (QIAGEN) according to the manufacturer’s protocol. RT-qPCR was used to measure RV NSP5 expression level as described previously (38).

### Statistical analysis

Bar graphs in **Fig. 2A-C** and **3A-B** were displayed as means ± standard deviation (SD). Bar graphs in **Fig. 4B, 5A-E, 6B, 7A-E, 8B-C, and S1** were displayed as means ± standard error of mean (SEM). Statistical significance in **Fig. 2A-C, 3A-B, 5A-E, 7A-E, and 8C** was analyzed by unpaired Student’s *t* test using GraphPad Prism 9.1.1. The asterisks in unpaired Student’s *t* test analyzed data represent * *P*<0.05; ** *P*<0.01; *** *P*<0.001; n.s., not significant. Statistical significance in **Fig. 4B, 6B, 8B, and S1** was calculated by Two-way ANOVA using GraphPad Prism 9.1.1. The asterisks in Two-way ANOVA analyzed data represent * *P*<0.05; ** *P*<0.01; *** *P*<0.001; n.s., not significant.

## ACKNOWLEDGEMENTS

The authors thank members of the Greenberg lab (Stanford University, USA) and Ding lab (Washington University in St. Louis) for constructive comments and suggestions. We very much appreciate Drs. Nathan J. Meade and Kenneth H. Mellits for kindly sharing the MA104-N*V cells and Dr. Phillippe H. Jais for sharing the C3P3 plasmid.

## REFERENCES

1. Dennehy PH. 2008. Rotavirus vaccines: an overview. Clin Microbiol Rev 21:198–208.

2. Crawford SE, Ramani S, Tate JE, Parashar UD, Svensson L, Hagbom M, Franco MA, Greenberg HB, O’Ryan M, Kang G, Desselberger U, Estes MK. 2017. Rotavirus infection. Nat Rev Dis Primers 3:17083.

3. Aliabadi N, Antoni S, Mwenda JM, Weldegebriel G, Biey JNM, Cheikh D, Fahmy K, Teleb N, Ashmony HA, Ahmed H, Daniels DS, Videbaek D, Wasley A, Singh S, de Oliveira LH, Rey-Benito G, Sanwogou NJ, Wijesinghe PR, Liyanage JBL, Nyambat B, Grabovac V, Heffelfinger JD, Fox K, Paladin FJ, Nakamura T, Agócs M, Murray J, Cherian T, Yen C, Parashar UD, Serhan F, Tate JE, Cohen AL. 2019. Global impact of rotavirus vaccine introduction on rotavirus hospitalisations among children under 5 years of age, 2008–16: findings from the Global Rotavirus Surveillance Network. The Lancet Global Health 7:e893–e903.

4. Troeger C, Khalil IA, Rao PC, Cao S, Blacker BF, Ahmed T, Armah G, Bines JE, Brewer TG, Colombara DV, Kang G, Kirkpatrick BD, Kirkwood CD, Mwenda JM, Parashar UD, Petri WA, Jr., Riddle MS, Steele AD, Thompson RL, Walson JL, Sanders JW, Mokdad AH, Murray CJL, Hay SI, Reiner RC, Jr. 2018. Rotavirus Vaccination and the Global Burden of Rotavirus Diarrhea Among Children Younger Than 5 Years. JAMA Pediatr 172:958–965.

5. Uygungil B, Bleesing JJ, Risma KA, McNeal MM, Rothenberg ME. 2010. Persistent rotavirus vaccine shedding in a new case of severe combined immunodeficiency: A reason to screen. Journal of Allergy and Clinical Immunology 125:270–271.

6. Velazquez FR. 2009. Protective effects of natural rotavirus infection. Pediatr Infect Dis J 28:S54–6.

7. Feng N, Yasukawa LL, Sen A, Greenberg HB. 2013. Permissive replication of homologous murine rotavirus in the mouse intestine is primarily regulated by VP4 and NSP1. J Virol 87:8307–16.

8. Sen A, Feng N, Ettayebi K, Hardy ME, Greenberg HB. 2009. IRF3 inhibition by rotavirus NSP1 is host cell and virus strain dependent but independent of NSP1 proteasomal degradation. J Virol 83:10322–35.

9. Barro M, Patton JT. 2005. Rotavirus nonstructural protein 1 subverts innate immune response by inducing degradation of IFN regulatory factor 3. Proc Natl Acad Sci U S A 102:4114–9.

10. Sen A, Rott L, Phan N, Mukherjee G, Greenberg HB. 2014. Rotavirus NSP1 protein inhibits interferon-mediated STAT1 activation. Journal of virology 88:41–53.

11. Graff JW, Ettayebi K, Hardy ME. 2009. Rotavirus NSP1 inhibits NFκB activation by inducing proteasome-dependent degradation of β-TrCP: a novel mechanism of IFN antagonism. PLoS Pathog 5:e1000280.

12. Arnold MM, Patton JT. 2011. Diversity of interferon antagonist activities mediated by NSP1 proteins of different rotavirus strains. J Virol 85:1970–9.

13. Graff JW, Mitzel DN, Weisend CM, Flenniken ML, Hardy ME. 2002. Interferon regulatory factor 3 is a cellular partner of rotavirus NSP1. J Virol 76:9545–50.

14. Arnold MM, Barro M, Patton JT. 2013. Rotavirus NSP1 mediates degradation of interferon regulatory factors through targeting of the dimerization domain. J Virol 87:9813–21.

15. Barro M, Patton JT. 2007. Rotavirus NSP1 inhibits expression of type I interferon by antagonizing the function of interferon regulatory factors IRF3, IRF5, and IRF7. J Virol 81:4473–81.

16. Lutz LM, Pace CR, Arnold MM. 2016. Rotavirus NSP1 Associates with Components of the Cullin RING Ligase Family of E3 Ubiquitin Ligases. J Virol 90:6036–48.

17. Ding S, Mooney N, Li B, Kelly MR, Feng N, Loktev AV, Sen A, Patton JT, Jackson PK, Greenberg HB. 2016. Comparative Proteomics Reveals Strain-Specific beta-TrCP Degradation via Rotavirus NSP1 Hijacking a Host Cullin-3-Rbx1 Complex. PLoS Pathog 12:e1005929.

18. Holloway G, Dang VT, Jans DA, Coulson BS. 2014. Rotavirus inhibits IFN-induced STAT nuclear translocation by a mechanism that acts after STAT binding to importin-alpha. J Gen Virol 95:1723–1733.

19. Kanai Y, Komoto S, Kawagishi T, Nouda R, Nagasawa N, Onishi M, Matsuura Y, Taniguchi K, Kobayashi T. 2017. Entirely plasmid-based reverse genetics system for rotaviruses. Proc Natl Acad Sci U S A 114:2349–2354.

20. Komoto S, Fukuda S, Ide T, Ito N, Sugiyama M, Yoshikawa T, Murata T, Taniguchi K. 2018. Generation of Recombinant Rotaviruses Expressing Fluorescent Proteins by Using an Optimized Reverse Genetics System. J Virol 92.

21. Feng N, Kim B, Fenaux M, Nguyen H, Vo P, Omary MB, Greenberg HB. 2008. Role of interferon in homologous and heterologous rotavirus infection in the intestines and extraintestinal organs of suckling mice. J Virol 82:7578–90.

22. Sanchez-Tacuba L, Feng N, Meade NJ, Mellits KH, Jais PH, Yasukawa LL, Resch TK, Jiang B, Lopez S, Ding S, Greenberg HB. 2020. An Optimized Reverse Genetics System Suitable for Efficient Recovery of Simian, Human, and Murine-Like Rotaviruses. J Virol 94.

23. Ding S, Zhu S, Ren L, Feng N, Song Y, Ge X, Li B, Flavell RA, Greenberg HB. 2018. Rotavirus VP3 targets MAVS for degradation to inhibit type III interferon expression in intestinal epithelial cells. Elife 7:e39494.

24. Patton JT, Taraporewala Z, Chen D, Chizhikov V, Jones M, Elhelu A, Collins M, Kearney K, Wagner M, Hoshino Y, Gouvea V. 2001. Effect of intragenic rearrangement and changes in the 3’ consensus sequence on NSP1 expression and rotavirus replication. J Virol 75:2076–86.

25. Zhao W, Xia M, Bridges-Malveo T, Cantu M, McNeal MM, Choi AH, Ward RL, Sestak K. 2005. Evaluation of rotavirus dsRNA load in specimens and body fluids from experimentally infected juvenile macaques by real-time PCR. Virology 341:248–56.

26. Pannacha P, Kanai Y, Kawagishi T, Nouda R, Nurdin JA, Yamasaki M, Nomura K, Lusiany T, Kobayashi T. 2021. Generation of recombinant rotaviruses encoding a split NanoLuc peptide tag. Biochem Biophys Res Commun 534:740–746.

27. Zhu S, Ding S, Wang P, Wei Z, Pan W, Palm NW, Yang Y, Yu H, Li HB, Wang G, Lei X, de Zoete MR, Zhao J, Zheng Y, Chen H, Zhao Y, Jurado KA, Feng N, Shan L, Kluger Y, Lu J, Abraham C, Fikrig E, Greenberg HB, Flavell RA. 2017. Nlrp9b inflammasome restricts rotavirus infection in intestinal epithelial cells. Nature 546:667–670.

28. Conceicao-Neto N, Zeller M, Lefrere H, De Bruyn P, Beller L, Deboutte W, Yinda CK, Lavigne R, Maes P, Van Ranst M, Heylen E, Matthijnssens J. 2015. Modular approach to customise sample preparation procedures for viral metagenomics: a reproducible protocol for virome analysis. Sci Rep 5:16532.

29. Yinda CK, Zeller M, Conceicao-Neto N, Maes P, Deboutte W, Beller L, Heylen E, Ghogomu SM, Van Ranst M, Matthijnssens J. 2016. Novel highly divergent reassortant bat rotaviruses in Cameroon, without evidence of zoonosis. Sci Rep 6:34209.

30. Bolger AM, Lohse M, Usadel B. 2014. Trimmomatic: a flexible trimmer for Illumina sequence data. Bioinformatics 30:2114–20.

31. Nurk S, Meleshko D, Korobeynikov A, Pevzner PA. 2017. metaSPAdes: a new versatile metagenomic assembler. Genome Res 27:824–834.

32. Buchfink B, Xie C, Huson DH. 2015. Fast and sensitive protein alignment using DIAMOND. Nature Methods 12:59–60.

33. Li H, Durbin R. 2010. Fast and accurate long-read alignment with Burrows-Wheeler transform. Bioinformatics 26:589–95.

34. Ali A, Roossinck MJ. 2007. Rapid and efficient purification of Cowpea chlorotic mottle virus by sucrose cushion ultracentrifugation. J Virol Methods 141:84–6.

35. Horie Y, Nakagomi O, Koshimura Y, Nakagomi T, Suzuki Y, Oka T, Sasaki S, Matsuda Y, Watanabe S. 1999. Diarrhea induction by rotavirus NSP4 in the homologous mouse model system. Virology 262:398–407.

36. Shaw RD, Hempson SJ, Mackow ER. 1995. Rotavirus diarrhea is caused by nonreplicating viral particles. Journal of virology 69:5946–5950.

37. Caddy SL, Vaysburd M, Wing M, Foss S, Andersen JT, O’Connell K, Mayes K, Higginson K, Iturriza-Gomara M, Desselberger U, James LC. 2020. Intracellular neutralisation of rotavirus by VP6-specific IgG. PLoS Pathog 16:e1008732.

38. Bolen CR, Ding S, Robek MD, Kleinstein SH. 2014. Dynamic expression profiling of type I and type III interferon-stimulated hepatocytes reveals a stable hierarchy of gene expression. Hepatology 59:1262–72.

